# Human mitochondrial variant annotation with HmtNote

**DOI:** 10.1101/600619

**Authors:** R. Preste, R. Clima, M. Attimonelli

## Abstract

HmtNote is a Python package to annotate human mitochondrial variants from VCF files.

Variants are annotated using a wide range of information, which are grouped into basic, cross-reference, variability and prediction subsets so that users can either select specific annotations of interest or use them altogether.

Annotations are performed using data from HmtVar, a recently published database of human mitochondrial variations, which collects information from several online resources as well as offering in-house pathogenicity predictions.

HmtNote also allows users to download a local annotation database, that can be used to annotate variants offline, without having to rely on an internet connection.

HmtNote is a free and open source package, and can be downloaded and installed from PyPI (https://pypi.org/project/hmtnote) or GitHub (https://github.com/robertopreste/HmtNote).

## Introduction

Recent Next Generation Sequencing (NGS) techniques allow researchers to collect an impressive amount of genomic information, particularly regarding genome variability. This is especially true for mitochondrial genomic research, which has seen a rapid increase in interest during the last few years, mostly due to the historically overlooked central role of the mitochondrion in many biological processes and pathological situations^1^. The presence of specific mitochondrial variants can offer useful insights on different levels, from population and evolution studies to pathogenicity assessment, and the functional analysis of human mitochondrial variations is a very florid research topic^2^.

Mitochondrial variants annotation, however, can be tricky to perform correctly, given the issue of mitochondrial heteroplasmy, a situation where a variable ratio of wild-type/variant mitochondrial genomes can be present in the same mitochondria, and the fact that some of the variants may represent benign variation only if related to the sample haplogroup; this may lead to uncertainty about which variants should be investigated further, and what effect they may exert in the living organism. Currently, a few tools for mtDNA variant annotation are available: MSeqDR mvTool^3^, SG-ADVISER mtDNA^4^, mtDNA-Server^5^, Mitomaster^6^, MitoTool^7^, Mitlmpact^8^; some of these resources, however, are not timely updated, and some others use a relatively narrow training set of mitochondrial genomes, so their annotations can be less reliable. A recently published resource, HmtVar^9^, offers variability and pathogenicity data about variants collected from over 45000 human mitochondrial genomes, thus rendering this information very useful to annotate variants found in a mitochondrial sample with a high confidence.

Here we propose a new tool for human mitochondrial variants annotation, HmtNote (https://github.com/robertopreste/HmtNote), which exploits the great amount of data available on HmtVar to enrich a given VCF file with functional annotations.

## Materials and methods

HmtNote aims at providing a simple yet efficient tool to annotate a list of human mitochondrial variants. It is built on Python 3 and functions as both an importable Python module and a command line interface (CLI) tool, so that it can easily be integrated into any NGS data analysis pipeline.

HmtNote operates on an input VCF file with human mitochondrial variants, which can be produced from any mitochondrial variant calling software; variants should be called with respect to the rCRS^10^, since HmtVar relies on this mitochondrial reference sequence. Every variant is looked for on HmtVar, exploiting its API, in order to retrieve all the available data. Annotations are distinguished into four annotation groups, which are detailed in Table 1, depending on the type of information they provide:

**Table 1.** Annotations provided by HmtNote.

| Annotation | Description | Annotation group |
| --- | --- | --- |
| Locus | Locus to which the variant belongs | Basic |
| AaChange | Aminoacidic change determined | Basic |
| Pathogenicity | Pathogenicity predicted by HmtVar | Basic |
| DiseaseScore | Disease score calculated by HmtVar | Basic |
| HmtVar | HmtVar ID of the variant | Basic |
| Clinvar | Clinvar ID of the variant | Cross-reference |
| dbSNP | dbSNP ID of the variant | Cross-reference |
| OMIM | OMIM ID of the variant | Cross-reference |
| MitomapAssociatedDiseases | Diseases associated to the variant according to Mitomap | Cross-reference |
| MitomapSomaticMutations | Diseases associated to the variant according to Mitomap Somatic Mutations | Cross-reference |
| NtVarH/<br>NtVarP | Nucleotide variability of the position in healthy/patient individuals | Variability |
| AaVarH/<br>AaVarP | Aminoacid variability of the position in healthy/patient individuals | Variability |
| AlleleFreqH/<br>AlleleFreqP | Allele frequency of the variant in healthy/patient individuals overall | Variability |
| AlleleFreqH_AF/<br>AlleleFreqP_AF | Allele frequency of the variant in healthy/patient individuals from Africa | Variability |
| AlleleFreqH_AM/<br>AlleleFreqP_AM | Allele frequency of the variant in healthy/patient individuals from America | Variability |
| AlleleFreqH_AS/<br>AlleleFreqP_AS | Allele frequency of the variant in healthy/patient individuals from Asia | Variability |
| AlleleFreqH_EU/<br>AlleleFreqP_EU | Allele frequency of the variant in healthy/patient individuals from Europe | Variability |
| AlleleFreqH_OC/<br>AlleleFreqP_OC | Allele frequency of the variant in healthy/patient individuals from Oceania | Variability |
| MutPred_Prediction/<br>MutPred_Probability | Pathogenicity prediction offered by MutPred and relative confidence | Predictions |
| Panther_Prediction/<br>Panther_Probability | Pathogenicity prediction offered by Panther and relative confidence | Predictions |
| PhDSNP_Prediction/<br>PhDSNP_Probability | Pathogenicity prediction offered by PhD SNP and relative confidence | Predictions |
| SNPsGO_Prediction/<br>SNPsGO_Probability | Pathogenicity prediction offered by SNPs & GO and relative confidence | Predictions |
| Polyphen2HumDiv_Prediction/<br>Polyphen2HumDiv_Probability | Pathogenicity prediction offered by Polyphen2 HumDiv and relative confidence | Predictions |
| Polyphen2HumVar_Prediction/<br>Polyphen2HumVar_Probability | Pathogenicity prediction offered by Polyphen2 HumVar and relative confidence | Predictions |

- basic data about a variant (locus, aminoacid change, pathogenicity, disease score, HmtVar ID);
- cross-reference information (Clinvar^11^ ID, dbSNP^12^ ID, OMIM^13^ ID, diseases associated to the variant according to Mitomap^6^);
- variability and allele frequency data (nucleotide and aminoacid variability in healthy and diseased individuals, allele frequency in healthy and diseased individuals from cumulative and continent-specific datasets);
- pathogenicity predictions from external resources (MutPred^14^, Panther^15^, PhD SNP^16^, SNPs & GO^17^, Polyphen2^18^).

Users can choose to use all of these groups of annotations, or select one or more specific annotation if preferred. HmtNote is able to efficiently annotate SNPs, insertions and deletions, as long as they are available on HmtVar.

HmtNote is best suited to work online, since it fetches annotations on-the-fly from HmtVar; nonetheless, it is also possible to use it even when no internet connection is available, thanks to its “offline mode”: users can download the required annotation database on their machine so that it can subsequently be used offline. The downloaded annotation database contains the very same information provided with online annotation, and details can be found in Table 1.

An extensive documentation about HmtNote and its usage and features can be found at http://hmtnote.readthedocs.io/.

## Results and discussion

In order to ensure that all annotations are correct and to provide users with some use cases, HmtNote was tested using a VCF file containing mitochondrial variants found in the 1KGenomes^19^ dataset. Annotations were performed using full annotation as well as specific basic, cross-reference, variability and prediction annotations options, both in online and offline mode to ensure that results were identical.

Out of the total 4242 variants reported in the VCF file, 3806 were annotated by HmtNote (more than 89% of the total number of variants). The detailed protocol used to download, process and annotate the VCF file is available in the related <u>project</u> on Open Science Framework^20^, together with the original and annotated VCF files.

## Conclusions

HmtNote is a simple and efficient tool to annotate human mitochondrial variants from VCF files. It exploits data hosted on HmtVar to perform its annotations using always the latest information available; in addition, it also allows to perform offline annotation if needed. Annotations can be limited to a specific type of information or span all the available data to provide a full enrichment of the annotated VCF file.

HmtNote employs an easy-to-use interface, which makes it appetible for both experienced bioinformaticians as well as scientists who are less familiar with programming, rendering the functional annotation of human mitochondrial variants a trivial task.

